# Are Genomic Language Models All You Need? Exploring Genomic Language Models on Protein Downstream Tasks

**DOI:** 10.1101/2024.05.20.594989

**Authors:** Sam Boshar, Evan Trop, Bernardo P. de Almeida, Liviu Copoiu, Thomas Pierrot

**Affiliations:** InstaDeep; Massachusetts Institute of Technology

## Abstract

Large language models, trained on enormous corpora of biological sequences, are state-of-the-art for downstream genomic and proteomic tasks. Since the genome contains the information to encode all proteins, genomic language models (gLMs) hold the potential to make downstream predictions not only about DNA sequences, but also about proteins. However, the performance of gLMs on protein tasks remains unknown, due to few tasks pairing proteins with the coding DNA sequences (CDS) that can be processed by gLMs. In this work, we curated five such datasets and used them to evaluate the performance of gLMs and proteomic language models (pLMs). We show that gLMs are competitive and even outperform their pLMs counterparts on some tasks. The best performance was achieved using the retrieved CDS compared to sampling strategies. We found that training a joint genomic-proteomic model outperforms each individual approach, showing that they capture different but complementary sequence representations, as we demonstrate through model interpretation of their embeddings. Lastly, we explored different genomic tokenization schemes to improve downstream protein performance. We trained a new Nucleotide Transformer (50M) foundation model with 3mer tokenization that outperforms its 6mer counterpart on protein tasks while maintaining performance on genomics tasks. The application of gLMs to proteomics offers the potential to leverage rich CDS data, and in the spirit of the central dogma, the possibility of a unified and synergistic approach to genomics and proteomics. We make our inference code, model weights and datasets available.

## Introduction

Large language models (LLMs), have revolutionized the field of NLP thanks to their ability to learn through self-supervision on unlabeled data [1–3]. Recently, the same techniques have been applied to learn from biological data. The discrete and sequential nature of biological sequences, such as proteins or DNA and RNA, paired with the abundance of unlabeled data, obtained through high-throughput sequencing, make it a perfect application for these methods to strive. This effort started first in proteomics, where several works showed that training large Transformer models to recover masked amino acids in protein sequences leads to powerful representations that can then be used to solve diverse downstream tasks with state-of-the-art performance [4–7]. More recently, similar models were developed for genomics and trained over the human reference genome as well as hundreds of reference genomes from different species to recover masked consecutive nucleotides in chunks [8–13]. These DNA models, while more recent and still less mature than their protein counterparts, have also showed the ability to build strong representations of nucleic acid sequences to solve downstream tasks with improved performance, including predicting diverse DNA molecular phenotypes related to splicing, regulatory elements and chromatin profiles.

Motivated by the central dogma of biology which states that the genome encodes all protein information, and by the fact that codon usage can influence protein structure and function [14], a third class of models, codon langauge models (cLMs), was recently introduced [15–17]. These models were trained on large datasets made of coding sequences (CDS) by reconstructing masked codons - rather than masked amino acids. Notably, the Codon Adaptation Language Model (CaLM) showed that cLMs can outperform their amino acid based counterparts on several downstream tasks of interest such as species recognition, prediction of protein and transcript abundance or melting point estimation when controlling for model size [15]. This improved performance seems to be related to the ability of codon-based language models to capture patterns of codon usage across DNA sequences.

Inspired by these recent results, we aimed to study to what extent genomic language models (gLMs) can be used as a general unified approach to solve tasks in both domains - genomics and proteomics. In opposition to cLMs, gLMs have been trained over full raw genomes and as such can be used to analyze non-coding regions as well as full genes including exons and intronic regions. While this makes gLMs widely capable for genomics tasks, their capacity to solve protein tasks from their corresponding CDS has not been explored. Since they have never seen "true" CDS per se during training, as exons are always separated by introns in eukaryotic species genomes, and coding sequences represent on average only ~ 1.5% of the human genome data used for training [18], it is unclear to what extent these models can be competitive with protein language models (pLMs).

In this paper, we present a comprehensive analysis of gLMs applied to protein-related tasks. We established a benchmark of five common protein analysis tasks and curated CDS sequences for a fair comparison between pLMs and gLMs. Our evaluation of two state-of-the-art pLMs and gLMs revealed that gLMs outperformed or matched pLMs on three out of five tasks, while underperforming on the remaining two. Notably, careful curation of matched CDS sequences was crucial for optimal gLM performance. The two tasks where pLMs significantly outperformed gLMs required sensitivity to codon-level changes. This observation led us to train a new gLM with a 3mer tokenization scheme, using full genomes as with our other gLMs. While our 3mer model slightly improved performance on the two codon-sensitive tasks, it did not enhance on the other proteins tasks and showed identical performance on genomic-specific tasks compared to its 6mer counterpart, suggesting that tokenization changes may not be the most fruitful path for improvement.

Intriguingly, gLMs significantly outperformed pLMs in predicting protein melting points – a trend also observed with cLMs. Further investigation revealed that gLMs achieve this by leveraging GC-content and species information from nucleotide sequences, features not captured by protein-based models. To better understand the representations learned by gLMs and pLMs, we designed a novel experimental protocol for systematically analyzing how these models represent sequences with single mutations within their embedding spaces. Interestingly, in the case of predicting beta-lactamase enzyme activity, gLMs clustered CDS by mutated positions, while pLMs grouped the corresponding proteins based on the mutated amino acid. These distinct representations motivated us to explore how to combine gLMs and pLMs to leverage their complementary strengths. We developed a new approach for combining both model classes. Our joint model recovers current state-of-the-art performance on four out of five tasks while setting a new state-of-the-art for protein stability prediction, demonstrating strong additive effects between their different sequence representations. We made the weights of our 3mer pre-trained model and the curated datasets available on HuggingFace.

## Results

### Genomic Language Models are competitive on protein tasks

In this work we studied the performance of gLMs (Fig. 1a) and pLMs (Fig. 1b) on five protein tasks of interest which are frequent in the literature [19, 20]. This collection includes sequence and residue level tasks, spanning regression and multi-label classification. Two of the tasks focus on a single protein and aim to predict the impact of a small number of mutations on the protein fitness, while the three other encompass different proteins coming from multiple topologies and species. We report the tasks along with the size of the respective datasets in Supplementary Table 1.

**Figure 1.**
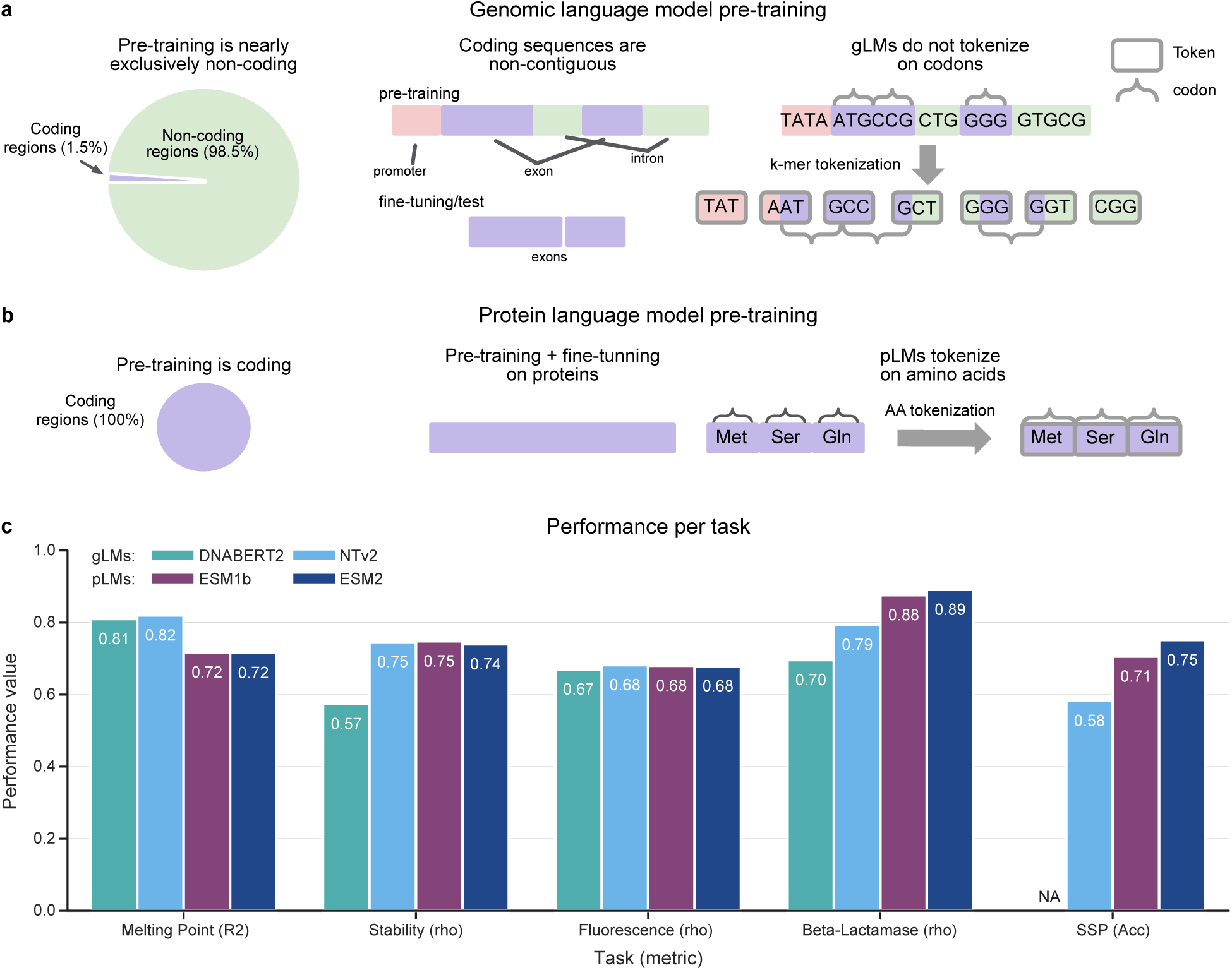
Genomic Language Models are Competitive on Protein Tasks. **a-b)** We outline key differences between gLMs and pLMs pre-training that make the task of building robust protein representations more difficult for gLMs. Unlike pLMs and cLMs, gLMs pre-training is predominantly (*∼* 99%) on non-coding regions of the genome, the vast majority of which (barring prokaryotic genomes) are non-contiguous, while fine-tuning and inference are carried out with contiguous CDS. Additionally, coding regions during pre-training are not tokenized on codons, making amino acid representations non-trivial. **c)** Evaluation results of Nucleotide Transformer v2 500M, DNABERT2, ESM2 650M, and ESM1-b 650M on the test datasets of the proposed tasks. The metrics used to measure performance were chosen to match previous benchmarks and include Spearman correlation *ρ*, *R*^2^, and accuracy, with a higher value indicating better performance for all metrics. Notably NTv2, matches or supersedes pLMs on 3 of the 5 tasks.

In order to be able to apply gLMs to protein tasks it is needed to create the DNA sequence-based version of the protein datasets, meaning the CDS sequence that encodes the proteins of interest (Fig. 1a). However, due to the degeneracy of the genetic code and the lack of protein identifiers in datasets, the process of obtaining the true CDS of a given protein sequence is not trivial. Here, we have retrieved, curated and released these associated CDS for each of the aforementioned tasks. We detail the exact process of retrieving these sequences in the Methods section. This dataset should enable further research into how gLMs may be applied to protein tasks.

For a comparison of pLMs with gLMs we decided to focus our attention on the ESM-2 [4] (650M parameters) and ESM-1b [21] (650M) models for proteins and NT-v2 [8] (500M) and DNABERT2 [9] (117M) for genomics, as these are considered as the state-of-the-art models in their respective fields. We evaluated these models by finetuning each on the five downstream protein tasks using the curated CDS sequences as inputs for gLMs and the respective amino acid sequences for pLMs (Fig. 1a,b). To ensure a fair comparison, we used the same methodology, number of steps and hyperparameters across all models. As recommended in recent work, we used a parameter-efficient finetuning approach [8]. See Methods for more details about our experimental protocol.

We compared the four aforementioned models over the five protein tasks (Fig. 1c). We excluded the secondary structure prediction task analysis for the DNABERT2 model as its tokenization technique yields to tokens representing variable numbers of nucleotides, thus making it challenging to link tokens to amino acids. For NT-v2 we simply made two predictions per embedding as the model uses a 6mer tokenization. First, we observed that the NT-v2 matches or outperforms its DNABERT2 gLM counterpart on all the protein downstream tasks, confirming the recently published results on genomics downstream tasks [8]. Interestingly, ESM-2 and ESM-1b seem to have comparable performance over these five tasks.

Our results show that NT-v2 matches or exceeds the performance of its pLMs counterparts on three of the five tasks: stability prediction, fluorescence prediction and melting point prediction (Fig. 1c). On melting point prediction in particular, NT-v2 and DNABERT2 models outperform significantly the ESM models. Interestingly, this result was also reported for cLMs [15]. We further explore these results later in this work. These results are noteworthy given the strong distribution shift between the distribution of the genomic data used to train gLMs and the distribution of curated CDS sequences. In particular, the vast majority of pre-training sequences are non-coding and the small fraction ( 1 2%) which are coding, are genes rather than CDS, and so coding exons are separated by non-coding introns. Furthermore, genomic tokenization yields tokens that, unlike proteomic or codon tokenization, are not identifiable with codons. These differences are summarized in Figures 1a and 1b. The results of Nucleotide Transformer on the protein tasks suggest that despite a significant distribution shift between pre-training and fine-tuning, as well as suboptimal protein tokenization, gLMs are able to capture protein features to the same extent than protein models.

However, we find that NT-v2 and DNABERT2 models underperform pLMs on the beta-lactamase activity prediction and secondary structure prediction tasks (Fig. 1c). This suggests that gLMs can capture global patterns in protein sequences but fail to capture finer-grained effects such as per amino acid structure or the impact of single point mutations, as assessed by secondary structure prediction and beta-lactamase respectively. We report here the results for the beta-lactamase *complete* dataset which contains all single codon subtiutions of the TEM-1 gene. We also explored whether synonymous codon information is improving gLM performance by evaluating all models on a maximal, non-degenerate, *unique* set and found lower performance (Supplementary Fig. 1). Results for the *unique* set as well as the procedure for generating both dataset variants is described in methods.

### True CDS sequences are essential to the performance of genomic language models

Since a growing body of biological literature indicates that codon bias influences protein folding [14, 22], there is a strong motivation to evaluate gLMs on the true CDS encoding the proteins of interest. However, this information is not always readily available, and when it is not, it can be a time-consuming, costly, or intractable process to retrieve the original CDS. As a result, in evaluating gLMs on proteins, it is common to approximate the desired CDS by sampling according to a given strategy. However, sampling from a global codon distribution may not recover biological plausible codon usage for a particular protein. Codon frequencies are known to vary not only between species (Supplementary Fig. 3), but also between genes of the same species [23], as we show for our five protein tasks (Fig. 2d,e, Supplementary Fig. 2). The implication is that it is unlikely that statistical sampling strategies can reconstruct the particular codon bias and relationships of a specific protein. The sampled CDS, although it may be drawn from some real codon distribution, may be unrealistic in the particular gene that it is sampled. While such studies have been conducting for codon language models, the effects of such imperfect codon sampling strategies on gLMs performance have not been studied yet.

**Figure 2.**
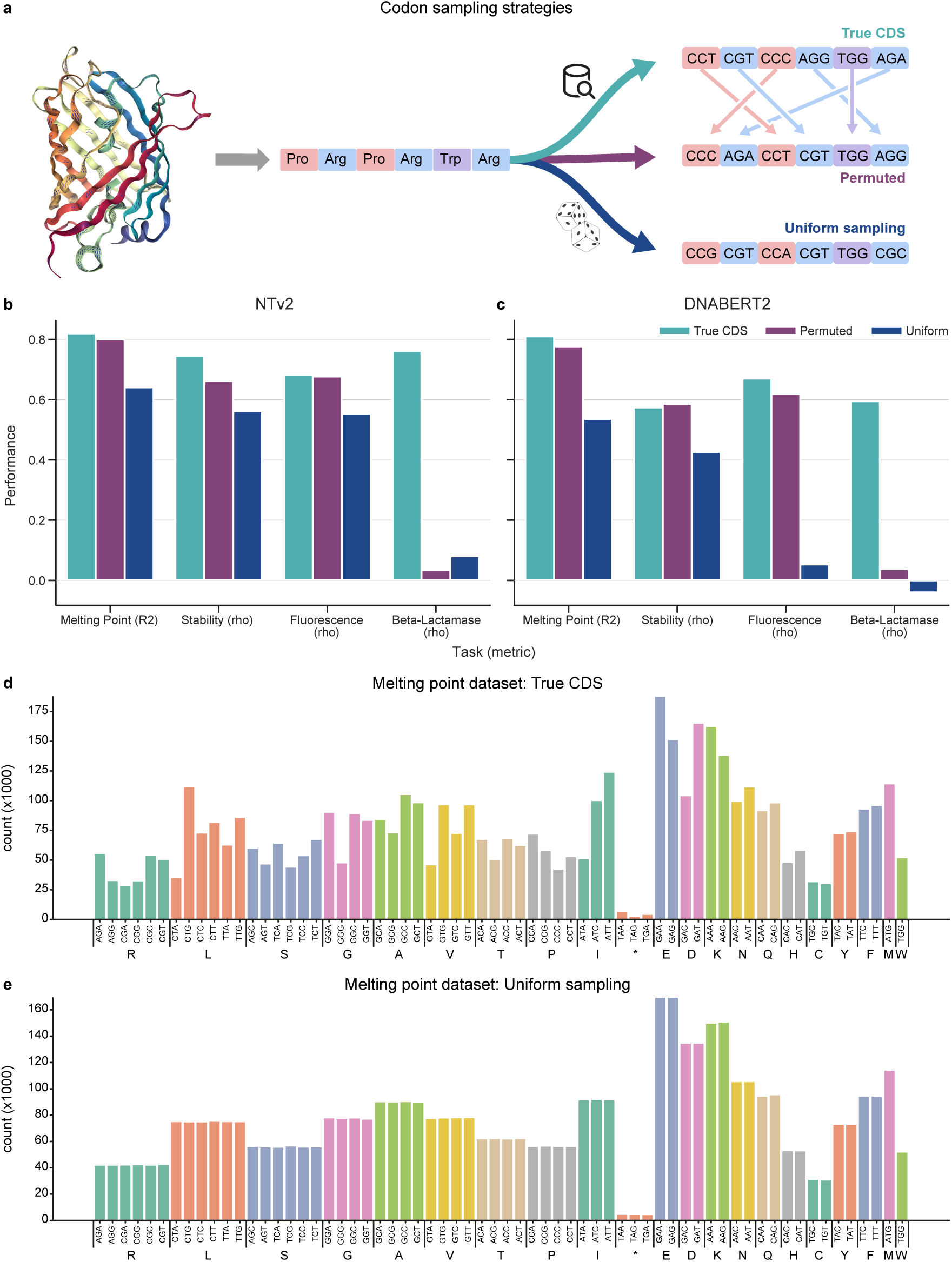
True CDS Outperforms Alternative Sampling Strategies. **a)** A schematic showing the three codon sampling strategies studied, all of which preserve the amino acid sequence. The uniform mutation describes uniformly sampling a codon from the set of its synonymous codons, the permutation mutation describe randomly shuffling codons of the same amino acid in the sequence, and the third using the true CDS. **b-c)** The impact of three codon sampling strategies on NT-v2 and DNABERT2 performance over 4 tasks. We leave the results for NT-v2 on SSP for the appendix since we could not evaluate DNABERT2 on SSP due to BPE tokenization. Performance is measured as Spearman correlation for Fluorescence, beta-lactamase, and Stability, *R*^2^ for Melting Point, and accuracy for SSP classification task. **d-e)** Disparity of codon frequencies between the true CDS distribution and uniform distribution for the melting point prediction task. We report the codon usage frequencies for each dataset in Supplementary Figure 2.

To investigate this question, we compared the performance of gLMs finetuned on the true CDS or on two other codon sampling strategies (Fig. 2a): (1) we permuted codons of like amino acids to perturb codon usage while respecting codon frequencies from the true CDS and (2) we sampled uniformly codons for each amino acid (note the difference in codon frequency between true and uniform sampling in Fig. 2d,e). We observed that on all tasks, having access to the "true" CDS improves the performance over sequences obtained by sampling codons from their natural frequencies (Fig. 2b,c), thus justifying the need for our curated dataset. We also show that randomly sampling codons yields degraded and close to zero performance on the beta-lactamase prediction task. These results were true for both NT-v2 and DNABERT2. In addition, we observed that NT-v2 is more robust than DNABERT2 to the codon distributions shift, which might be related to the different tokenizations, 6mers tokenization and byte-pair encoding (BPE), respectively.

### 3mer tokenization is not enough to close the gap between gLMs and pLMs

Given that pLMs were only superior to gLMs on the two protein tasks that required fine-grained amino acid-level resolution (beta-lactamase and Secondary Structure Prediction; Fig. 1c), and as codons are represented by groups of 3 nucleotides, we investigated whether 3mer tokenization in NT-v2 could be more suitable for those tasks than the current 6mer tokenization scheme. To address this, we pre-trained a 50M parameter NT-v2 model but replacing the 6mer tokenization by a 3mer one, while respecting the exact same training data (full genomes) and hyperparameters than NT-v2 models [8] (Fig. 3a, Supplementary Fig. 4a,b). We then compared the performance of this newly pre-trained model to its 6mer counterpart as well as to ESM-2 150M, the closest

**Figure 3.**
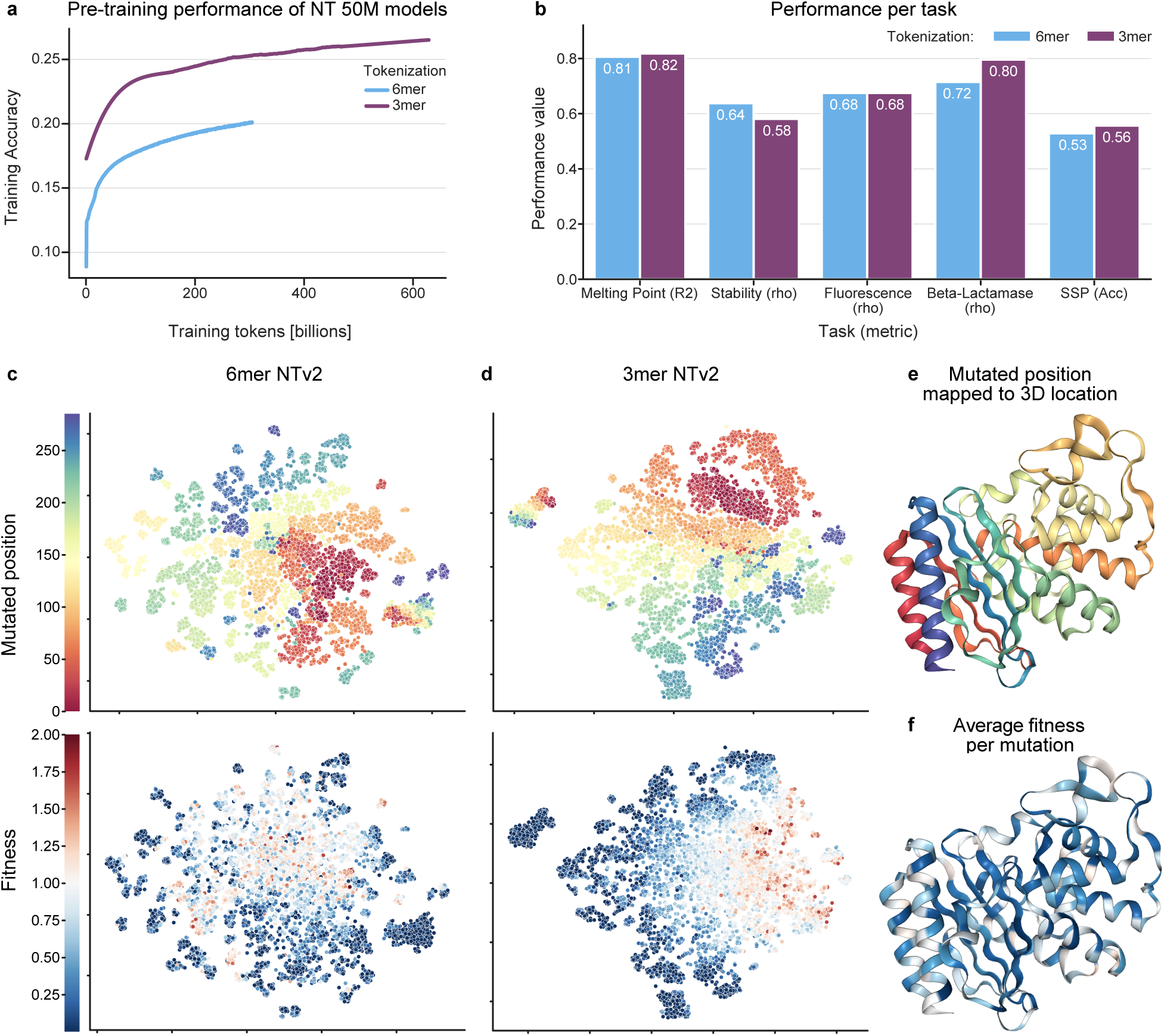
3mer Tokenization Achieves Better Performance than 6-mer on Specific Protein tasks. **a)** Pre-training validation accuracy of the 3-mer and 6-mer model on 600B tokens and 300B tokens respectively. **b)** Test set performance of 3-mer model at the 300B checkpoint and 6-mer model at the 300B checkpoint. The 3-mer tokenized model outperforms the 6-mer on all tasks but one, despite seeing half of the nucleotide base pairs in pre-training. **c-d)** Dimensionality reduction via t-SNE on the training set of the beta-lactamase enzyme activity task reveals that the 3-mer model better organizes sequences in relation to fitness and organizes similarly to the 6-mer model in regards to mutated position. Here we take WT to be fitness 1.0 and color it white. Fitness values above WT are colored red, and those below are colored blue. This result may suggest that the more fine-grained 3-mer resolution allows NT-v2 to better predict the single codon changes that compromise this dataset. **e)** Mapping mutated position to the 3D structure of the protein reveals clustering by secondary structure. **f)** Fitness averaged over synonymous mutation shows that most variants perform worse than WT.

ESM-2 model in terms of number of parameters.

We observed that the 3mer tokenization improves performance over 6mer tokenization on the beta-lactamase activity and structure prediction tasks (Fig. 3b). These two tasks require codon-level precision, which can explain why models that tokenize codons individually may have an advantage. However, this improved performance was not enough to close the gap with pLMs. In addition, 3mer tokenization does not improve over two other tasks and even decreases performance on the third task.

We note that as a result of coarser tokenization, the 6mer model has seen twice as many nucleotide base pairs in training as the 3mer model. For completeness, we trained the 3mer model up to the same number of base pairs (600 billion tokens). We find that despite consistent improvement measured by the pre-training objective, we see little change in performance on the protein downstream tasks (Supplementary Fig. 4c and Supplementary Table 3).

A t-SNE dimensionality reduction of beta-lactamse embeddings of the finetuned 3mer and 6mer models shows that the 3mer model organizes better the embeddings with respect to sequence fitness, supporting the evaluation results and the hypothesis that fine-grained tokenization is valuable for isolating the effects of single codon mutations (Fig.3c-e). We also note that under the reduction, both models cluster embeddings of sequences which are mutated at similar locations in the pimrary structure of the protein (Fig.3c-e). However, the 3mer model seems to globally organizes embeddings by position mutated, while the 6mer model only locally clusters sequences with similar mutated positions.

We also evaluated the 3mer tokenization model on the 18 downstream genomics tasks from the NT study [8] to evaluate the impact of the tokenization on genomics datasets. Here we observed identical performance, within the margin of error, with its 6mer counterpart (Supplementary Fig.4d,e). Grouping more nucleotides per token is beneficial to gLMs as it allows to extend their perception field, showed to improve performance [24, 25], while keeping the number of tokens constant and hence preventing the quadratic scaling of compute of Transformer models. As such, 6mer tokenization reduces compute time and cost compared to 3mer tokenization. As the 3mer tokenization does not yield significant overall improvement on protein downstream tasks, only on the ones that require codon-level precision, 6mer tokenization is to be preferred for gLMs. This also suggests that changing the tokenization scheme of gLMs might not be the most fruitful path towards advancing these models.

### gLMs dominate for melting point prediction through the identification of GC-content, species and codon usage

We showed that gLMs outperforms significantly their pLM counterpart on the melting point prediction task. A similar behavior has been reported for cLMs [15]. This motivated us to analyze the disparity between gLMs and pLMs performance on this task. In particular, we explored whether the superior performance of gLMs can be attributed to a biological phenomenon such as codon usage, or whether it is exploiting a “superficial” feature unique to CDS data. Here we define superficial as information readily available that does not contribute to a better understanding of proteins.

In investigating the impact of codon usage reported in Figure 2a,b, we found that in the absence of codon usage information the NT-v2 performance drops below that of ESM, the gLM achieving an accuracy of only 0.64 compared to the pLM’s 0.72 (Fig. 1c). This result suggests that NT-v2 is utilizing codon frequencies. We next explored if the improved performance on the melting point prediction task could be related to additional sequence features. One indication that the NT-v2 might be exploiting superficial features of CDS would be if it can achieve the similar performance using only global sequence information. The motivation is that a biological phenomenon regarding codon usage would likely depend on their absolute and relative locations. To test this we developed two hypotheses around the use of global sequence information.

### The GC-content Hypothesis

We hypothesized that the NT-v2 may use GC-content to influence protein melting point prediction. The GC-content of a genomic sequence indicates the proportion of guanine (G) or cytosine (C) bases. G-C base pairs, featuring three hydrogen bonds, are more stable than A-T base pairs with two hydrogen bonds. Higher GC-content leads to higher melting temperatures in equal-length sequences. To test this hypothesis, we augmented both ESM-2 and NT-v2 with the sequence’s GC-content information by appending the normalized GC-content to the embeddings before making the melting point prediction. Although this addition moderately improves performance with an increase in *R*^2^ from 0.72 to 0.74, the model still lags behind NT-v2 (Fig. 4a). Augmenting NT-v2 with the same information does not lead to any increase in performance. This suggests that NT-v2 already has access to GC-content information.

**Figure 4.**
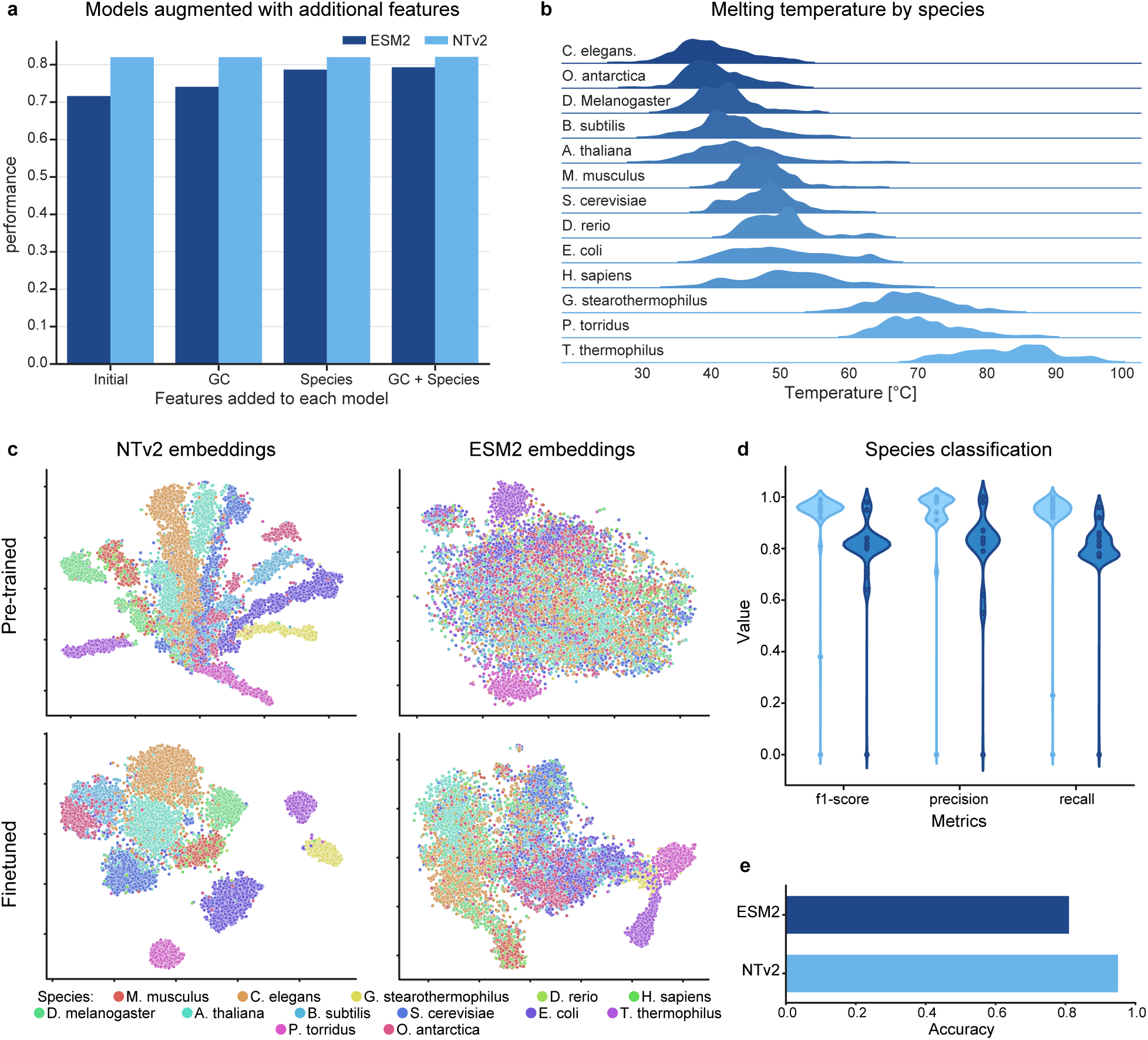
gLMs Use Species Codon Usage Bias to Outperform pLMs on Melting Point Prediction. **a)** The results of appending combinations of GC-content and Species to NT-v2 and ESM-2 embeddings during fine-tuning on the melting point prediction task. We find that species information accounts for the majority of the disparity of performance between ESM-2 and NT-v2. We also augment NT with the same information but see no change in performance indicating NT-v2 already has access to this information. We report full results in Supplementary Table 5. **b)** The distribution of melting points for each species in the dataset show distinct profiles. **c)** Dimensionality reduction via t-SNE of the pre-trained and fine-tuned NT-v2 and ESM-2 models demonstrates that the gLM captures the structure of species information to a greater degree than pLM and initially acquired this knowledge from its pre-training. **d)** We train ESM-2 and NT-v2 models to predict the species from sequence via fine-tuning with a single layer classification head. We plot the f1-score, precision and recall across species, weighted by the number of sequences in each species. **e)** Bar plot for the overall species classification accuracy. Results from both d and e confirm that NT is superior at identifying species. We present full confusion matrices in Supplementary Figures 5 and 6.

### The Species-level Conditioning Hypothesis

We next explored if the NT-v2 may exploit codon usage information to condition on the species the sequence was derived from. The melting point prediction dataset consists of proteins from thirteen different species ranging from unicellular E. coli, to mice and humans. Proteins of different species have distinct melting point profiles and identifiable codon preferences (Fig. 4b). Codon bias across species is a well-documented phenomenon that reflects mutational and selective pressures [26], and is evident in the melting point prediction dataset (Supplementary Fig. 3). To test this hypothesis, first we verify that gLM can better identify species from sequence. We finetuned NT-v2 and ESM-2 on the task of species identification and found that NT achieves an accuracy of 0.95 while ESM-2 achieves an accuracy of only 0.81 (Fig. 4d,e; Supplementary Fig. 5, 6). Additionally, we showed that the t-SNE for pretrained embeddings of models reveal that gLM embeddings are strongly structured by species while pLM are not (Fig. 4c).

To test whether species information may account for the difference in performance we augmented both ESM-2 and NT-v2 with the species information of each sequence and evaluated test set performance. This augmentation was done by appending a one-hot species-identifying vector to the embeddings of each model. We find that augmenting ESM-2 with species information increases performance from an *R*^2^ value of 0.72 to 0.79 (Fig. 4a). This closes most of the gap with the NT-v2 trained from curated CDS and brings the model to the performance of NT-v2 trained with permutated codons (no local information) which has an *R*^2^ of 0.80. In contrast, augmenting NT-v2 with species does not result in an improvement in performance (Supplementary Table 5), suggesting that NT-v2 achieves the majority of its advantage via conditioning on species information, which it learned during pre-training.

Using these findings, we finally tested if augmenting ESM-2 with both global attributes (GC-content and species) could recover the performance of NT-v2. Our results show that although there are additive benefits for ESM-2 from having both features, the majority of the information appears to come from species identification, and there still exists a gap in performance with NT-v2 (Fig. 4a, Supplementary Table 5). We presume this remaining advantage is coming from local codon interaction information present in the coding sequences.

### gLMs and pLMs organize local embedding structure by different principles

Following the melting point prediction analysis and the different performances between gLMs and pLMs, we aimed to better understand the differences in sequence representation between the different model types. To do so, we systematically visualized through t-SNEs the embeddings spaces of finetuned NT-v2 and ESM-2 models on all tasks. We have previously observed clustering by species in the emebdding space of NT-v2 for the melting point prediction task (Fig. 4c). We also observed clustering by mutated positions and amino acids at mutated positions for models finetuned on the beta-lactamase activity prediction task (Fig. 5d,e). In order to confirm numerically this finding, we derived a novel experimental protocol to investigate local structure in the embedding space of each type of model (Fig. 5a). Our protocol aims to determine whether mutated sequences whose embeddings are in the same neighborhood in the finetuned model latent space, share similar properties with one another. To do so, we determined for each sequence in the dataset the k-nearest neighbors in each finetuned model latent space, using mean embeddings over tokens, and computed similarity metrics over the sequences properties. In the case of beta-lactamase, we looked specifically at the distance between mutated positions as well as similarities of amino acids in the mutated sequence that we named here amino acid accuracy.

**Figure 5.**
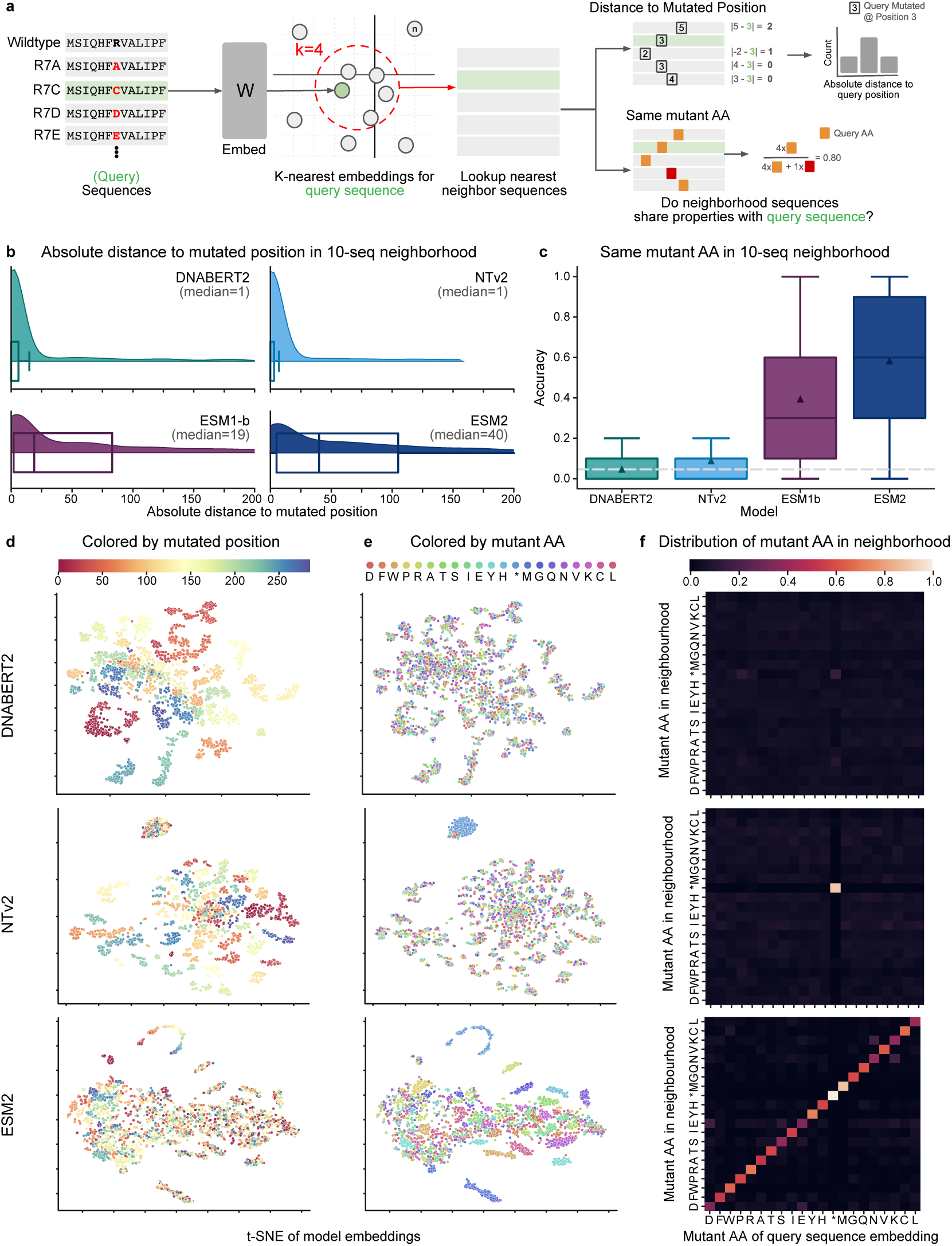
gLM and pLM Organize Protein Embeddings via Different Sequence Properties. **a)** A schematic depicting the experimental procedure used to probe the model embedding spaces. The beta-lactamase enzyme activity task consists of all single codon mutations of the TEM1 gene, however we limit ourselves to a maximal non-degenerate subset. The procedure first embeds sequences, searches for nearest neighbors using euclidean distance, and then determines whether neighbors share certain properties with the query sequence. **b)** Distributions of the absolute difference between the mutated position of the query sequence and that of its 10 sequence neighborhood. Results show that the median absolute distance between the position of the query sequence and its neighbors is much larger for pLMs than gLMs. **c)** Boxplot of the percent of sequences in each 10 sequence neighborhood which share the same mutated amino acid. The dotted line indicates the expected accuracy under a uniformly random distribution of embeddings, while the white triangles indicate the mean accuracy. **d)** t-SNE of model embeddings averaged across the sequence length and colored by the location of the mutation. **e)** The same t-SNE of model embeddings but now colored by the mutant amino acid. **f)** Heatmaps where the x axis denotes the mutant amino acid of the query sequence and the column is the distribution of mutated amino acids of sequences whose embeddings are in the 10-neighborhood of that query vector.

Our method demonstrates that gLMs locally organize their sequences by the location of the mutation (Fig. 5d,e,f). In particular, we find that the median distance between the mutated position of a sequence and its 10 nearest neighbors is only 1 for gLMs, while for pLMs it is significantly larger (Fig. 5b). We report results for a neighborhood of size 10, but found that the results held for larger neighborhoods (Supplementary Fig. 7a).

Unlike gLMs, we observed that finetuned pLMs organize their sequences by the identity of the mutated amino acid (Fig. 5d,e,f). That is, sequences embeddings tend to be close if the mutant codon translates to the same amino acid. We find that for ESM-2 and ESM-1b, in a 10-embedding neighborhood sequences share the same mutant amino acid as the query sequences with accuracy 58.4% and 39.4% respectively (Fig. 5c). This is significantly above the expected accuracy of 4.6% where the embeddings arrange uniformly randomly with respect to mutant amino acid identity (Fig. 5c). The NT-v2 and DNABERT2 models have an amino acid accuracy of 8.8% and 4.8% respectively. Similarly to the mutated position, we find that this analysis is robust with respect to neighborhood size (Supplementary Fig. 7b).

### Genomic and Proteomic Language Models are Additive

Finally, motivated by the competitive, and in some cases superior, performance of gLMs on our benchmark, as well as by the differences observed in gLMs and pLMs sequence latent spaces, we studied whether the two approaches may be complementary for solving protein tasks. To this end, we finetuned and evaluated a joint genomic-protein language model across all regression tasks. The joint model consists of NT-v2 and ESM-2 models with their final embedding concatenated together followed by a single layer regression head (Fig. 6a). The NT-v2 component uses the true CDS while the ESM-2 component uses the respective translated protein. The joint model is trained identically to each of its sub-models using identical training strategy and hyperparameters.

**Figure 6.**
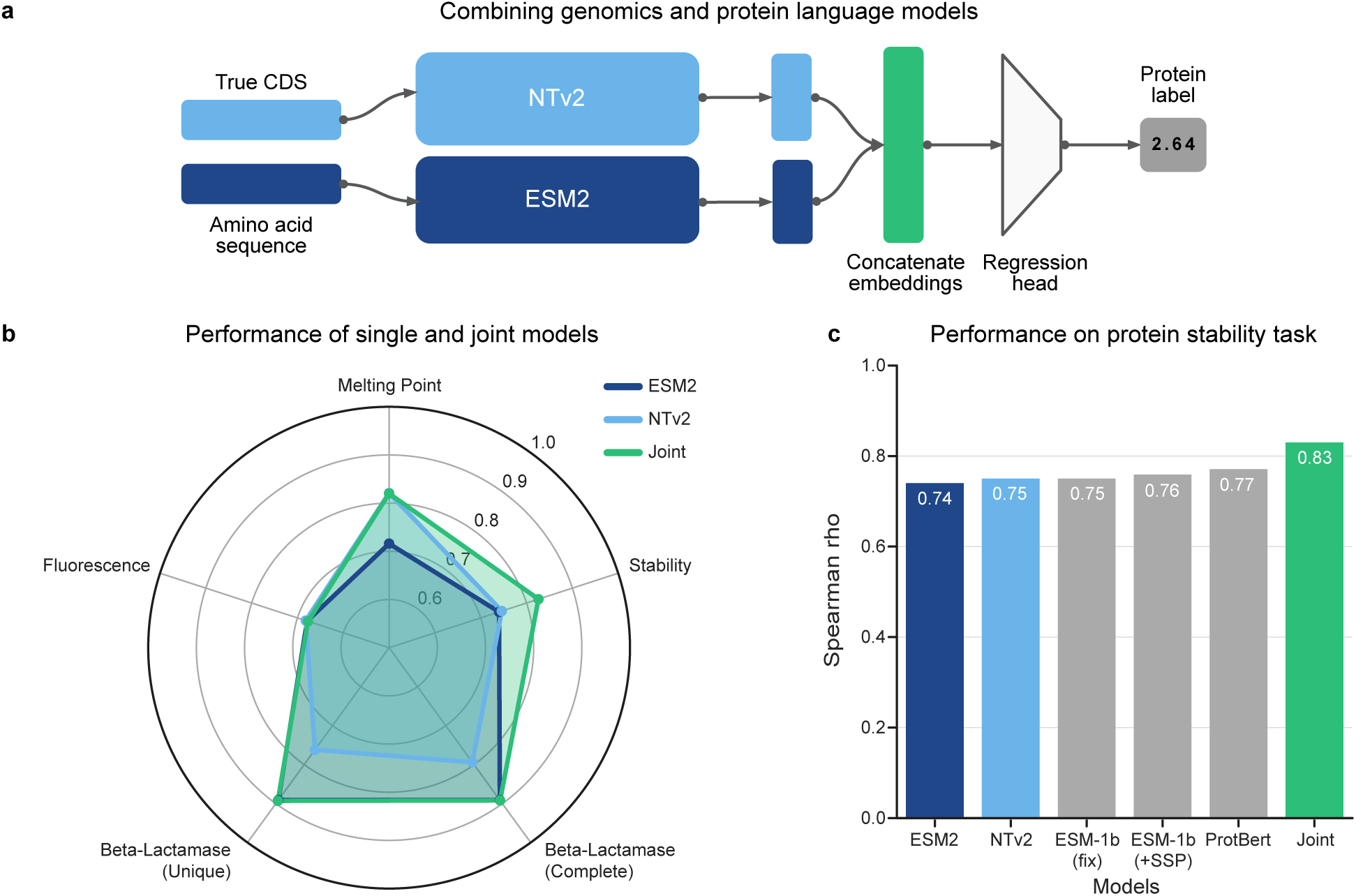
Genomic and proteomic language models are additive. **a)** Schematic depicting how the joint model was constructed. The true CDS and amino acid sequences were given as input to NT-v2 and ESM-2 respectively. Output embeddings from each model were averaged over the sequence length, concatenated together, and fed to a predictor head. **b)** Radar plot of the test set performance on the four regression tasks proposed in the work. Results show that the joint model matches over outperforms its single model counterparts. **c)** Spearman correlation on the test set for the protein stability task exhibits that the joint genomic-proteomic outperforms state of the art models from literature.

We found that on all tasks the joint model performs at least as well as genomic and proteomic sub-models (Fig. 6b). In particular, on the task of the stability prediction, the joint model outperforms each individual approach and achieves a new state-of-the art performance against the PEER Benchmark for Protein Sequence Understanding [19], even outperforming approaches which utilize additional external data and multi-task learning (Fig. 6c). Overall, our results show that each type of model captures different but complementary sequence representations that can be combined in joint genomic-proteomic models to achieve improved performance on protein tasks. The success of this joint genomic-proteomic approaches motivates continued research into how we might build protein models that leverage the best of both approaches.

## Discussion

To the best of our knowledge, this work represents the first attempt to evaluate the potential of gLMs to also solve protein tasks. In order to do that, we retrieve, curate, consolidate and publish protein datasets with the respective CDS sequences that encode these proteins. We show that evaluating gLMs on true CDS is the fairest way to compare them to pLM on protein tasks, and offers gLM the potential to leverage codon information that may influence protein structure [14, 22]. We show that gLMs perform consistently better on true CDS than on sequences generated from other, even quite similar, sampling strategies. We release the datasets publicly with the hope that they will encourage further research to build general models that can be widely used across fields in biology including genomics and proteomics.

We evaluated two state-of-the-art gLMs (NT-v2 & DNABERT2) and two pLMs (ESM-2 & ESM-1b) across this benchmark, offering the first preliminary comparison of the two approaches on protein downstream tasks. We demonstrate that gLMs are surprisingly capable on protein downstream tasks, especially given that their pre-training on whole genomes and non-contiguous coding regions are not conducive to learning protein representations. In particular, we find that gLMs match or exceed pLMs on 3 of the 5 tasks of interest, even exceeding pLM performance on melting point prediction.

On melting point prediction, we find that gLMs greatly outperform their protein counterparts, as previously reported for the cLM CaLM [15]. We investigate this disparity in performance and explore whether such models represent better protein sequences or whether they capture other superficial feature of the CDS not present in the translated amino acid sequence. We find that it is in large part explainable by gLMs’ ability to identify and condition on the species of a sequence based on the codon bias fingerprint present in the CDS.

We note that NT, and gLMs in general, underperform relative to pLMs on the two tasks that require codon-level fine-grained precision. We investigate whether this performance difference may, in part, be due to the NTs coarser 6mer tokenization. To this end we pre-trained from scratch a 3mer tokenized NT-v2 model and compared it to an identically trained and sized 6mer model, finding that the finer tokenization consistently performs better on those two downstream protein tasks. This suggests that while the 6mer tokenization may be an ideal trade-off for gLMs on genomic tasks, which often benefit from longer context windows, for protein tasks the coarseness of multi-residue tokenization may come at a cost.

In addition, we explored whether gLMs and pLMs represent proteins in a fundamentally different way. We offer some preliminary analysis that focuses on how each model type organizes its sequence embedding space. Our results suggest that the two methods structure local neighborhoods of embeddings according to different principles. In particular, local regions of protein embedding space shared common mutated amino acids, while local regions of genomic embedding spaces shared a common location of mutation in the primary structure of the protein. These differences in structure exist in the embeddings space, but continue to be apparent when the dimensionality is reduced.

Since gLMs achieved state-of-the-art performance on the majority of the protein downstream tasks and capture different sequence representations than pLMs, we explored whether gLMs and pLMs may be additive. Indeed, we found that each type of model captured different but complementary sequence representations and that a joint genomic-proteomic model supersedes both individual approaches. This results suggests that the synergy of gLMs and pLMs may be a promising future direction for improving the performance on protein tasks.

The demonstrated successes of gLMs in modeling protein sequences indicate that they might be a good starting point to build unified foundational models for biology, but there is still much work needed to understand how to improve these models. Our findings suggest that gLMs and pLMs have different strengths and represent proteins in different ways. This observation not only further motivates research on the interpretability of gLMs for protein tasks, but also on joint methods that may extract the advantages of both approaches. We expect that the collection and release of the five CDS datasets will help the community to make progress in this direction.

## Methods

### Models

Language models (LMs) are a statistical method for modeling language. These methods create a probability distribution over words, giving the likelihood that a sequence of words exists. A common way to train LM is with so-called cloze tests where the models is tasked with predicting a masked word (or words) given the context of the sentence. This is known as masked language modeling (MLM). Masked language modeling has seen incredible adoption an success because it allows one to leverage enormous amounts of unlabeled data to learn good representations. Models trained in an unsupervised manner with the masked language objective can then be fine-tuned for particular tasks.

This same approach has been widely adopted in biology where there is a large amount of unlabeled sequence data and relatively few labels. These sequences can be treated much like sentences. First applied to proteins, this approach has now seen success in modeling genomic sequences. The most powerful and common architecture for language modeling is the transformer. The transformer architecture operates on sets of tokens which are vector embeddings of components of the input sequences. If the input us an image, a token could be the vector of pixels in a patch. If the input is a sentence, the token might be a vector embedding of a sequence of characters, or a word. Indeed, this approach has seen success in both natural language processing (NLP) and computer vision.

When using the MLM objective to train language models, the most common approach is to use Bidrectional Encoder Representations from Transformers (BERT) [1]. This approach allows the language model to condition on the entire context of a masked token in pre-training, rather than perhaps just the left-context which as approach commonly used in auto-regressive human language modeling. Since biological sequences do not have the left-to-right structure of human sentences, it integral to utilize bidrectional information. Embeddings representations of masked tokens from BERT are mapped to a probability distribution over the token vocabulary by a language modeling head.

### Architecture

All models considered in this work were encoder-only transformers. The genomic language models considered were closely related to the ESM family of architectures [4], allowing a fair comparison between the genomic and proteomic approaches. All architectures share some main components. The first is an encoding layers which transforms tokens into an embedding space. The second is a transformer stack, which accepts this numerical representation and is trained to refine token representations in the context of the surrounding sequence. All models also have a head which transformers the refined embedding to some desired output. In pre-training models utilize a language model head which takes the final embedding representations of the model and converts it into a probability distribution over the vocabulary of tokens. In fine-tuning, the language model head is ignored and a single-layer task-specific head responsible for transforming these final representations into predictions. As we are primarily studying existing pre-trained models and their fine-tuned performance and embeddings, we are predominantly concerned with the second setting.

### Transformer Block

The transformer block is the key component of the ESM architecture. Each transformer component consists of layer normalization followed by multi-headed self-attention. Self-attention is the mechanism that is central to the transformer architecture and allows all tokens to attend to one another in data-dependent dynamic manner. This is implemented via scaled dot product attention.

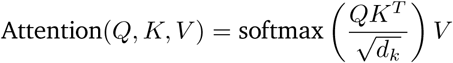

Here each token submits a query and key vectors of dimension *d*_*k*_, as well as a value vector. Each are stacked in matrices *Q*, *K* and *V* respectively. The dot product of the query matrix *Q* and the key matrix *K*, calculates the similarity between each pair of query and key vectors. These attention values are normalized to a probability distribution with softmax and then used to dynamically scale the set of value vectors, *V* . Intuitively, the process asks how much any two tokens should attend to one another (the attention value), and does a weighted sum of the tokens value vectors according to the calculated attention.

### Tokenization and Token Embedding

Although these models are quite similar, they do differ in the way in which they tokenize an input sequence, and the process by which they encode these tokens into a numerical representation.

#### ESM

Both ESM architecture tokenize on amino acids which is a natural way to break up protein sequences. The ESM1b model uses learned positional embeddings to embed tokens. The ESM2 architecture utilizes Rotary Position Embeddings (RoPE) [27] which, unlike learned embeddings, allow the extrapolation to longer sequences than trained on.

#### Nucleotide Transformer v2

The Nucleotide Transformer v2 uses 6mer tokenization on DNA sequences. This coarser tokenization allows the model to accept sequences as long as 12kbp, which is particularly important for capturing long range genomic interactions. As described in the Nucleotide Transformer paper [8], the second version of the model, similar to ESM, makes use of the RoPE embeddings scheme.

#### DNABert2

DNABert2 uses the Byte Pair Encoding scheme, rather than k-mers to tokenize DNA sequences [9]. BPE is an method first developed for encoding strings of text which iterative combines the most frequently occurring n-gram of length 2. The authors argue that this approach is more computationally efficient. DNABert2 also replaces the learned positional embeddings of its predecessor with Attention with Linear Biases (ALiBi) [28], an method which, similar to RoPE, removes the constraint on input length.

#### Pre-training

We pre-trained the 3mer 50 million parameter Nucleotide Transformer identically to its 6mer counterpart described in the original Nucleotide Transformer paper [8]. The model was trained on 64 TPUv4s for 8 days on the Multispecies dataset (described in detail in the same paper [8]). We used batch size of 8 sequences and a fixed length of 2048 tokens. Sampled sequences had 15% of their tokens altered. Of this subset 80% were masked. We added noise by replacing an additional 15% of tokens with randomly selected standard tokens.

We utilized the batch sum of the cross-entropy between the predicted probabilities and the target tokens as the loss. The effective batch size was chosen to be 512, corresponding to 512 2048 1million tokens. We used Adam optimizer [29] and learning rate scheduling as described in the Nucleotide Transformer paper [8], with *β*_1_ = 0.9, *β*_2_ = 0.999 and *ε* = 1*e* 8. Over the first 16,000 steps, we utilized a warmup schedule which increased the learning rate linearly from 5*e* 5 to 1*e* 4, and then decreasing as the squareroot decay over the rest of pre-training. Validation was performed on a heldout set every 5*e*9 tokens, and checkpoints were saved every 1*e*10 tokens.

The 3mer model was trained until 600 billion tokens to match the number of base pairs which the 6mer model (trained to 300 billion tokens) had seen. However, we report the 3mer results for the checkpoint at 300B tokens as we did not see a significant increase in performance on the downstream tasks between the two checkpoints (Supplementary Table 3).

#### Fine-tuning

Fine-tuning of the models was done using IA^3^ [30] parameter-efficient fine-tuning, along with a single-layer classification or regression head. IA^3^ scales activation by a learnable vector, introducing a number of parameters approximately 0.1% of the total number of parameters. Models were fine-tuned with a batch size of 8. Adam optimizer was used with a learning rate of 0.003. Models were evaluated at fixed intervals over the validation set during training. Checkpoints with the highest *R*^2^ for regression and lowest cross-entropy loss for classification over the validation set were saved and evaluated on the test set. For a given task all models were trained with the same number of steps which we report in Supplementary Table 4.

#### Evaluation Methodology

The two pre-trained gLMs, DNABERT2 and NT-v2, and the two pre-trained pLMs, ESM1b and ESM2, were respectively evaluated with corresponding CDS and protein sequences as input and fine-tuned in similar conditions for a fair comparison. In opposition to all the other tasks that are regression tasks at the sequence level, the SSP task is a classification task at the amino acid level. This is simply performed by pLMs by predicting for each amino acid embedding a secondary structure from the 8 possible classes (Supplementary Table 6). For the Nucleotide Transformer, as tokens represent 6-mers, each token embedding is mapped to two classification predictions corresponding to the two amino acids that the 6-mer represents. As DNABert2 uses Byte Pair Encoding to tokenize nucleotides sequences, we couldn’t retrieve any systematic mapping from tokens to amino acids and thus couldn’t evaluate this model over the SSP task.

### Protein Downstream Tasks Datasets

We study five protein tasks of interest that are frequent in the literature. This collection includes sequence- and residue-level tasks, spanning regression and multi-label classification. We detail and motivate below these five tasks. See Table 1 for an overview of these tasks.

### Tasks

#### Secondary Structure Prediction (SSP)

Understanding the structure of proteins is integral to understanding their function. This task tests a model’s ability to learn local secondary structure. The task is a multi-label classification task where each input amino acid is associated with one of 8 labels, denoting which secondary structure that residue is a part of. All secondary structures were empirically derived using crystallography or NMR.

The structural data for the training and validation sets were collected by Klausen *et al* [31]. Crystal structures were retrieved from Protein Data Bank and filtered with a 25% sequence similarity threshold, a resolution at least as fine as 2.5 angstrom and a length of at least 20 amino acids [31]. Following the work of Klausen [32] we used splits filtered at 25% sequence identity to ensure generalization, and evaluated on 3 test sets: CASP12, CB513, TS115.

Models were evaluated on three independent test datasets: TS115 [33], CB513 [34] and CASP12 [35]. TS115 consists of 115 proteins whose structure has been determined by X-ray crystallography, filtered to a resolution of 3 Angstroms [33]. CB513 is composed of 513 non-redundant protein regions from 434 proteins whose structures are determined by X-ray crystallography [34]. CASP12 is a curated list of 21 proteins whose structures have been determined through crystallography or NMR [35].

#### Melting Point Prediction (MPP)

It is often desirable to design protein with specific thermostability profiles. However, predicting protein melting point can be a challenging task as even single residue mutations can have large impacts [36]. Melting point prediction (MPP) is a sequence-level regression task that evaluates a model’s ability to predict a measure of melting temperature.

The data orginates from the thermostability atlas, which was originally measured and compiled using a mass spectrometry-based proteomic approach [37]. We follow the same “mixed” splits described in FLIP [38] which seek to avoid over-emphasis of large clusters. Sequences are clustered at 20% identity with 80% of clusters assigned to the train dataset and 20% of clusters assigned to the test dataset.

#### Beta-lactamase Activity Prediction

It is also important for models to have the precision to accurately predict the effects of single amino acid mutations [19]. Beta-Lactamase is a regression task consisting of sequences from a study exploring the fitness landscape of all single codon substitutions in the TEM-1 gene [39]. Labels indicate the ability of mutant genes to confer ampicillin resistance.

The data for this task is from Firnberg *et al* [39] which systematically examined fitness landscape of all single codon mutations in the TEM-1 Beta-lacatamase gene synthesized in native host E. coli. The TEM-1 gene is known to confer antibiotic resistance, and fitness is taken to be a function of this resistance. In particular, gene fitness was measured by splitting the library of mutants onto thirteen sub-libraries exposed to increasing levels of ampicillin concentration. Using deep sequencing, the resistant alleles were counted in each sub-library. Unnormalized fitness *f*_*i*_ of allele *i* was calculated as a weighted average of the allele counts on each plate by the log Amp concentration [39]. Fitness values were then normalized by wildtype fitness, *f*_*WT*_ , such that normalized fitness *w*_*i*_ > 1 indicates a mutation which is more fit that wildtype [39].

Since beta-lactamase task consists of all single codon mutations, the dataset contains many degenerate coding sequences. In PEER [19] labels were averaged over degenerate coding sequences in the original dataset. This process removes much data and does not allow us to study gLMs on degenerate sequences, as well as the impact of synonymous mutation on fitness. Consequently, we propose two training datasets, sharing a single test set. The *Complete* set contains all CDS samples except those that are degenerate with respect to any CDS in the test set. The *Unique* set contains a random, maximal, subset of the non-degenerate coding sequences. This Unique set allows comparison between the gLMs and pLMs since all translated sequences are unique, while the Complete set demonstrates the impact of data availability on gLM performance. Notably, although the unique dataset is non-degenerate we use the raw fitness values of the CDS, rather than those averaged over degenerate sequences.

#### Fluorescence Prediction

Estimating the fitness landscape of proteins which are many mutations away from the wildtype sequence is one of the core challenges of protein design. This task evaluates a model’s ability to predict log-fluorescence of higher-order mutant green fluorescent protein (avGFP) sequences.

Original data is from an experimental study of the avGFP fitness landscape [40]. The library was generated via random mutagensis of the wildtype sequence and synthesized in E. coli. Inspired from the TAPE and PEER benchmarks [19, 20], we restrict the training set to amino acid sequences with three or fewer mutations from parent GFP sequences, while the test set is all sequences with four or more mutations.

Like the Beta-lactamase task, random mutagenesis of the avGFP gene led to CDS which were degenerate. However, since this process was much less systematic, this was true of a much smaller fraction of the sequences. In the training set and validation sets there were 54,025 uniquely translating CDS of the 58,417 total sequences. Since most sequences were non-degenerate we selected a random maximal subset and did not study the degenerate sequences. Notably, since the test set was higher order mutants, there was only one amino acid sequence (SS26C:SN168H:SD188V:SS200G) of the 27,217 which did not have a unique coding sequence. We removed randomly one of the two corresponding CDS. Notably, this means the test set is near identical to that used in TAPE [20] and PEER [19] benchmarks.

#### Protein Stability Prediction

It is important for models trained on diverse sequences to be able to accurately predict a small region of the fitness landscape. This regression task evaluates how well models predict stability around a small region of high-fitness sequences. The train and validation originate from a multi-round experiment and consist of a broad selection of de novo computationally designed proteins composing a small number of topologies. The test set consists of the neighborhoods of single codon mutations around a few of the most stable candidates [41].

The data for this task originates from Rocklin *et al* [41] in which stability is measured as a function of the resistance to increasing levels of protease. In particular, the designed libraries were synthesized in yeast and exposed to different concentrations of protease. At each level of protease the fraction of proteins remaining folded was measured, and these values were used to infer the *EC*_50_: the value at which half of cells express proteins that pass a defined stability threshold. The stability of a protein is then defined as the difference between the *EC*_50_ value of the protein and that of the predicted *EC*_50_ in the unfolded state, calculated in a log_10_ scale [41].

#### Performance Metrics

Since we have chosen tasks frequent in the literature, we follow the standard performance metrics that have been used in order to be able to compare our methods to previous results. For Beta-Lactamase Activity , Fluorescence and Stability we use Spearman’s *ρ*. For Melting Point Prediction we use the coefficient of determination (*R*^2^), and for Secondary Structure Prediction we use per amino acid accuracy.

### Retrieving and Curating Coding Sequences

One main contribution of this work is to retrieve, curate, and share consolidated CDS datasets for the five protein tasks of interest to allow the comparison of nucleic acid- and amino acid-based models. We detail in this paragraph how these CDS were collected for each task.

#### Melting Point Prediction

For MPP, we used the Uniprot[42] ID mapping tool to map the Uniprot ID’s associated with each protein, available from the TAPE benchmark [20], to the DNA sequence database of EMBL CDS [43]. Any retrieved CDS from EMBL whose translation did not match the original amino acid sequences were filtered out.

#### Secondary Structure Prediction

In SSP, we used protein sequences with associated PDB ID’s [44] from the dataset hosted by Netsurf P-3.0 [32]. To collect the CDS we first used the RCSB 1D Coordinate Server [44] which assembles alignments between structure and sequence databases, to find alignments to protein sequences from the Uniprot database. Returned alignments to Uniprot were filtered out if there was not complete coverage. The remaining Uniprot id’s were then mapped to the sequence database EMBL CDS using the same process as for MPP described above.

#### Beta-Lactamase Activity Prediction

For the beta-lactamase task, all sequences corresponded to the same gene. We obtained the TEM-1 reference gene as well as the mutations from supplementary material of ref. [41].

#### Stability Prediction

For the stability prediction task, coding sequences were taken from supplementary material of the original experimental study [41]. Since all CDS translate into unique amino acids, we are able to match the dataset splits presented in TAPE [20].

#### Fluorescence Prediction

Finally, for the fluorescence task we obtained the reference GFP gene, as well as its mutations from the reference of the original data [40]. We chose to take the *Unique* subset as described above since the dataset was mostly non-degenerate.

### Additional Analysis

#### Nearest Neighbor Analysis

We define the *k* neighborhood of a sequence *s* to be the set of sequences whose embeddings are the *k* nearest neighbors of *s*’s embedding. More formally, let *f*_θ_ : *V* ^*m*^ → *R*^*d*^ be a model mapping sequences in *V* ^*m*^ to some *d*-dimensional embedding space. Here *V* is the token vocabulary of the model, *m* the fixed length of the input, and θ the model’s parameterization.

We can define the *k*-nearest neighbors of *s*_*i*_ *∈ S* with respect some model *f*_θ_, to be the set *NN*_*k*,θ_(*s*_*i*_) *⊂ S* such that |*NN*_*k*,θ_(*s*_*i*_)| = *k* and *∀s*_*j*_ *∈ S ∖ NN*_*k*,θ_(*s*_*i*_)

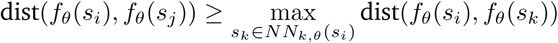

where dist is some metric on the embeddings space. In this work we take this metric to be the Euclidean distance. We also experiment with cosine similarity and find similar results. It is important to note that cosine similarity is not truly a distance metric as it doesn’t satisfy the triangle inqeuality. There are also some concerns regarding fidelity and reliability of cosine similarity in high dimensions [45]. For these reasons we choose to use the Euclidean distance.

#### TSNE Dimensionality Reduction

For the study of model embedding spaces, we chose to use TSNE to reduce the embedding dimensions so that they could be visualized in two dimensions. Although we experimented with other dimensionality reduction methods, we chosen TSNE because our study focused on local neighborhoods of embeddings and TSNE primarily maintains the relative distances between neighboring points. A future study looking at global organizing principles of embeddings spaces may choose to use another reducing method that respects global relationships between points (eg UMAP).

#### A Comment on the meaning of "true CDS"

In this work we use True CDS both to mean the transcript which really encoded a protein of interest, but also to mean a natural transcript that could have encoded the protein of interest. It is not always possible to retreive the coding sequence which truly encoded the measured protein, as the study of interest may not have recorded in it. In the cases in which the coding sequence were recorded (Fluorescence, Beta-Lactamase, Stability) we were able evaluate the genomic language models on these sequences.

In the cases where the coding sequences were not available (Secondary Structure, Melting Point), we followed the aforementioned procedure detailed in methods. That is, we retrieved a biological transcript, present in a real reference genome, which encoded the protein of interest. While this doesn’t guarantee that the transcript is identical to that which truly produced the protein, we argue that it the closest you can recover, and most importantly, we know with certainty that it is biological plausible. Research has shown that codon bias can vary within genes of the same species, and that certain codon may be important at specific locations in a gene. Sampling strategies generally do not respect these codon relationships, while our method does.

## Data Availability

HuggingFace versions of our CDS-protein task datasets can be found at https://huggingface.co/datasets/ InstaDeepAI/true-cds-protein-tasks.

## Code Availability

Model code and weights of the 3mer pre-trained transformer model as well as inference code in Jax are available for research purposes at https://github.com/instadeepai/nucleotide-transformer. HuggingFace version of the model, in PyTorch, can be found at https://huggingface.co/InstaDeepAI/ nucleotide-transformer-v2-50m-3mer-multi-species.

## Acknowledgements

We would like to thank Patrick Bordes for help on assembling the CDS dataset for the MPP task.

## Supplementary Figures

**Supplementary Figure 1.**
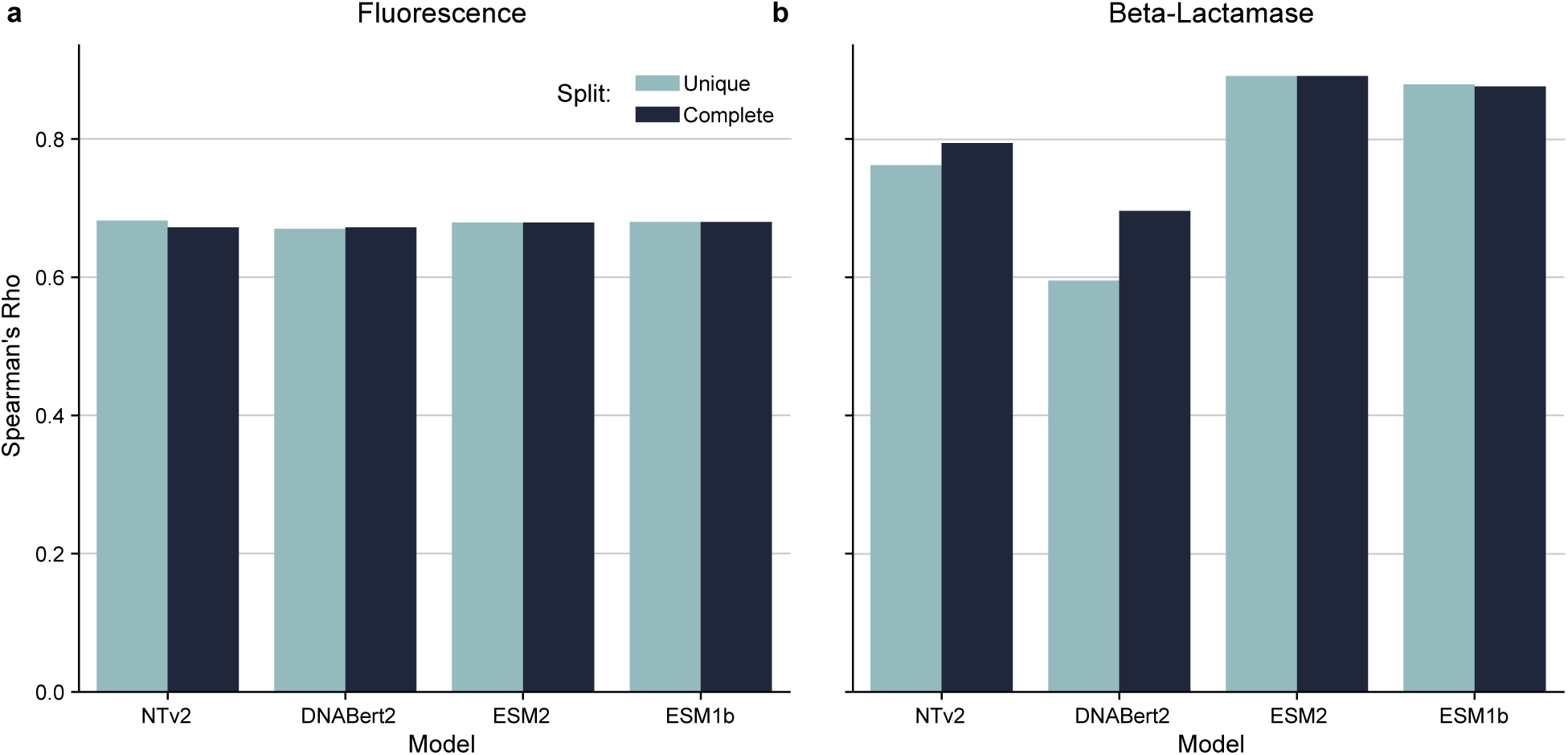
Training on Synonymous Codon Improves gLM Beta-Lactamase Performance. **a)** Model performances is nearly identical between Fluorescence Complete and Unique set. This is because, as described in Methods , there are hardly any degenerate sequences. **b)** Performance of models on Beta-Lactamase Complete vs Unique datasets, evaluated on the same test set. GLMs trained on the complete dataset outperform those trained on the Unique Set. Unlike Fluroescence there are many more degenerate sequences in the Complete Set. There is no observed effect for pLMs.

**Supplementary Figure 2.**
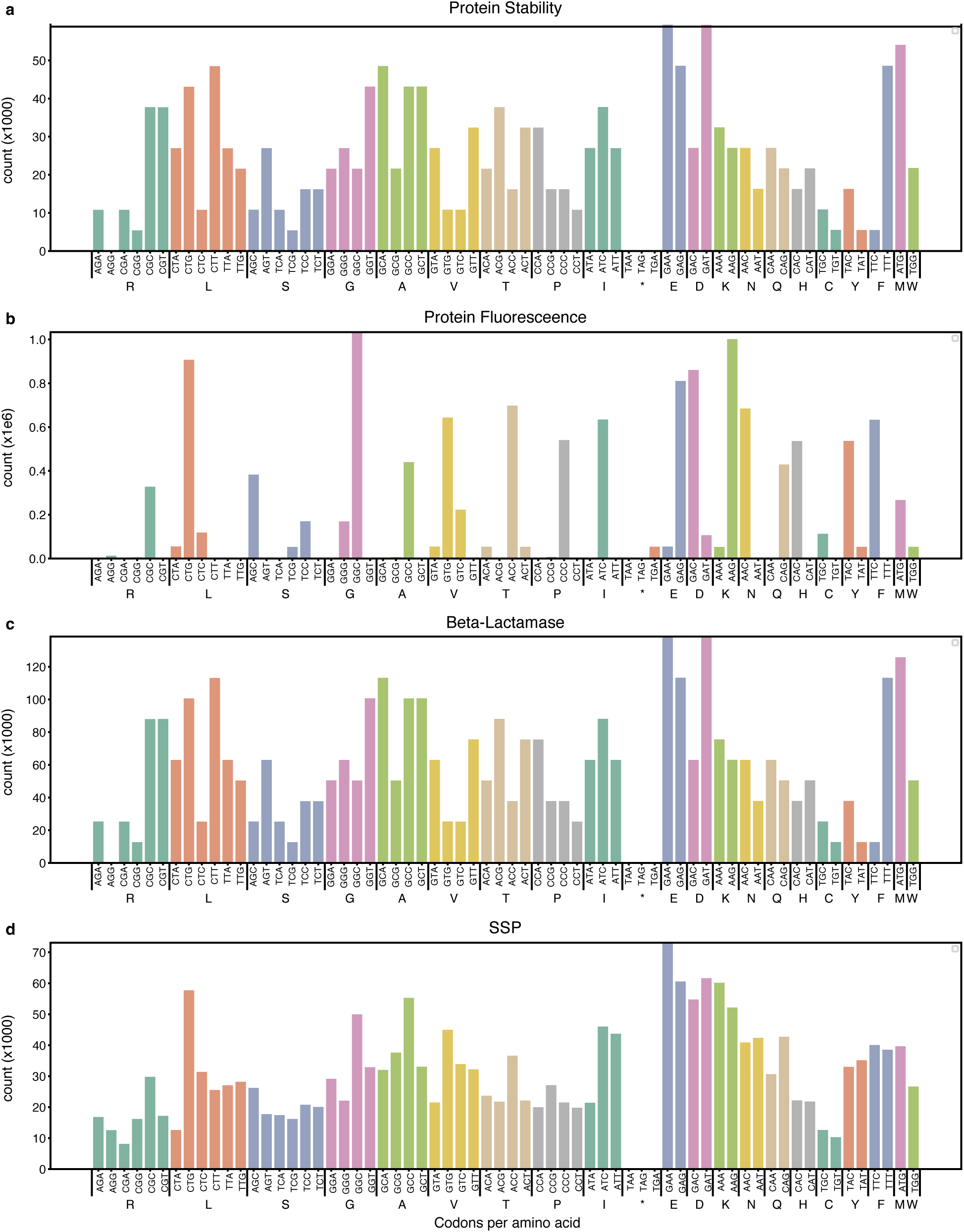
Codon frequency for the proteins present in each dataset.

**Supplementary Figure 3.**
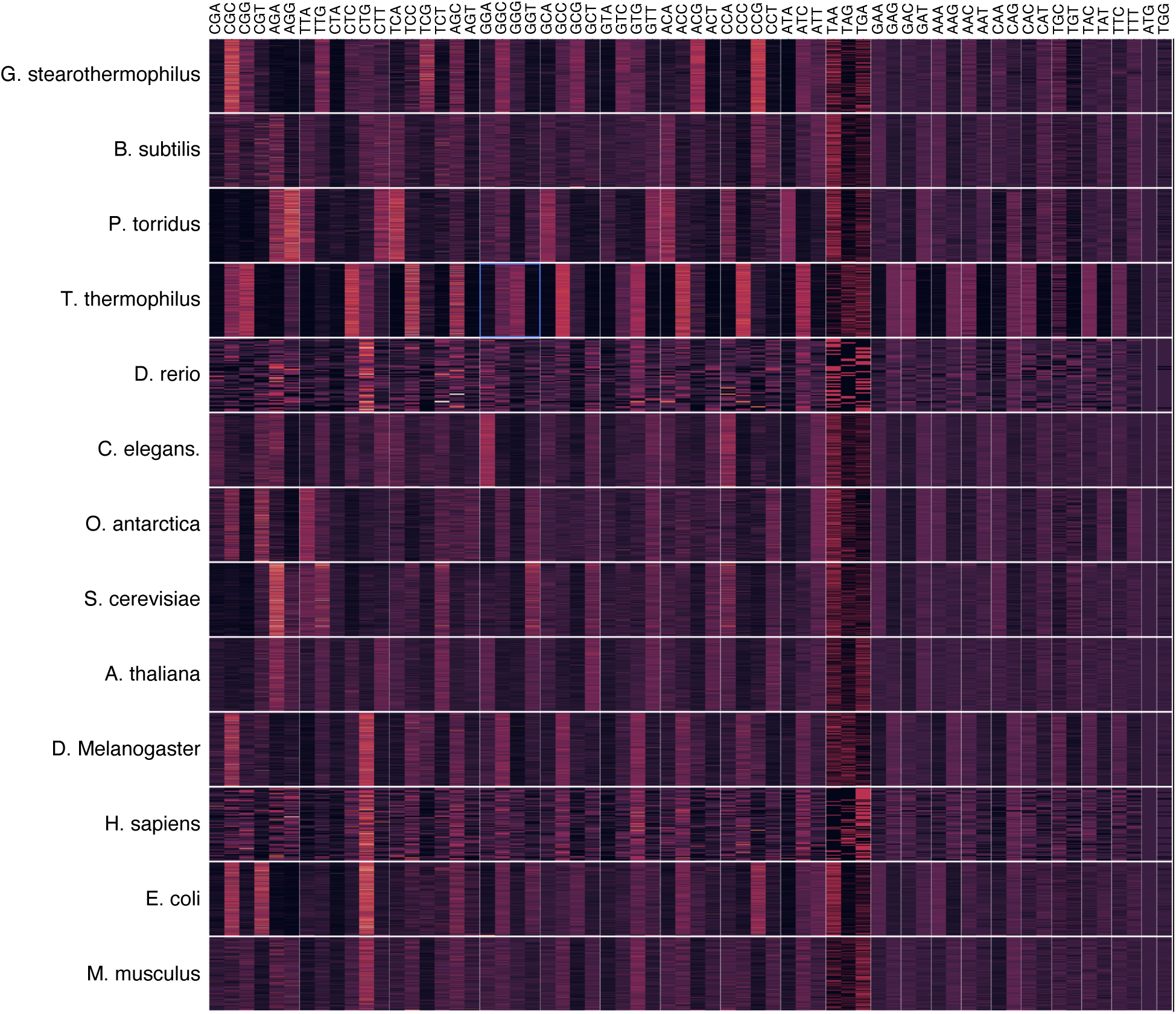
Relative synonymous codon usage per species. This figure shows the relative synonymous codon usage (RSCU) of each sequence in the melting point task, partitioned by species. RCSU is defined as the observed codon frequency over the expected codon frequency assuming synonymous codons are equally likely. In the figure, columns are amino acids and sub-columns are codons. Rows are species, and sub-rows are individual sequences with that species. Thus, the color at some sub-row, sub-col location denotes the RCSU of some codon in a a sequence of a particular species, where bright colors are closer to 1. The figure reveals many species have strong and identifiable synonymous codon preferences.

**Supplementary Figure 4.**
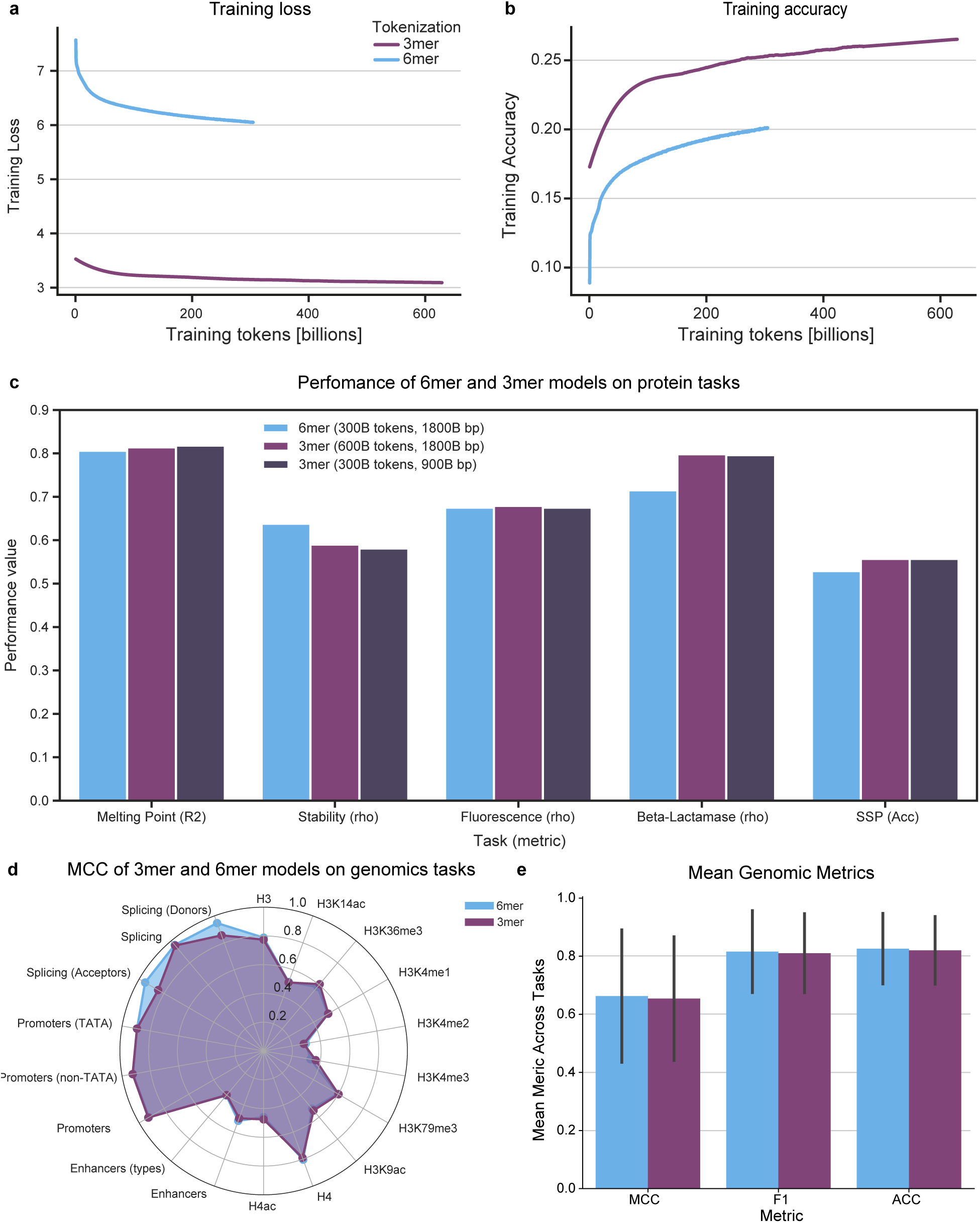
Improved performance of 3mer over 6mer NTv2 on protein tasks. **a-b)** Training curves for the pre-training of the 3mer and 6mer NTv2 models. Training loss **(a)** and accuracy **(b)** are shown in function of the number of training tokens (billions). **c)** Performance of the 6mer and two checkpoints of the 3mer model per task. The metric used for each task is mentioned in the x-axis labels. **d)** Matthew’s correlation coefficient (MCC) for 3mer (300B tokens) vs 6mer tokenized models across 18 genomics tasks. **e)** Mean MCC, F1-score and Accuracy across all genomics tasks for 3mer (300B tokens) and 6mer models.

**Supplementary Figure 5.**
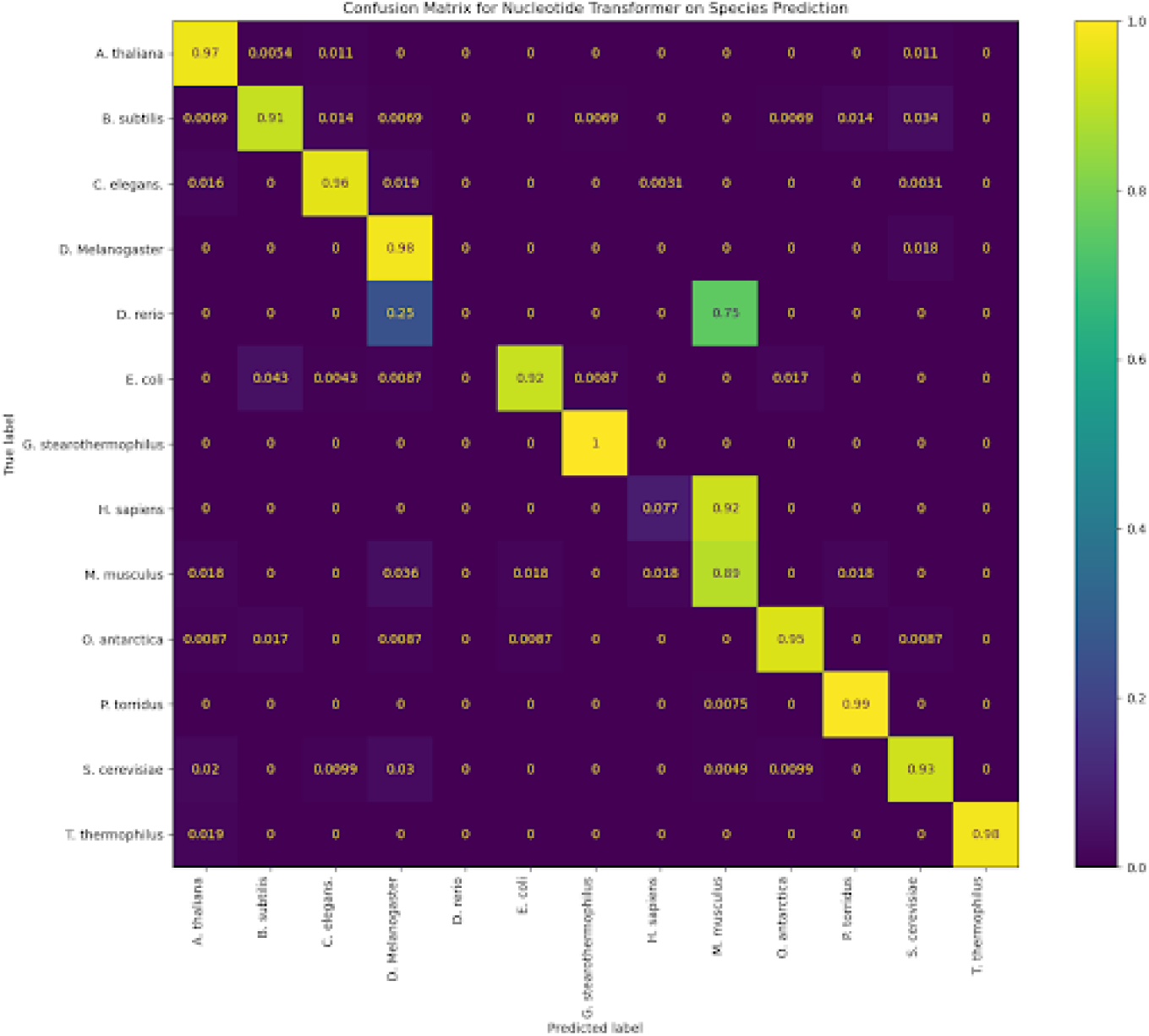
Confusion matrix for NTv2 model on species prediction.

**Supplementary Figure 6.**
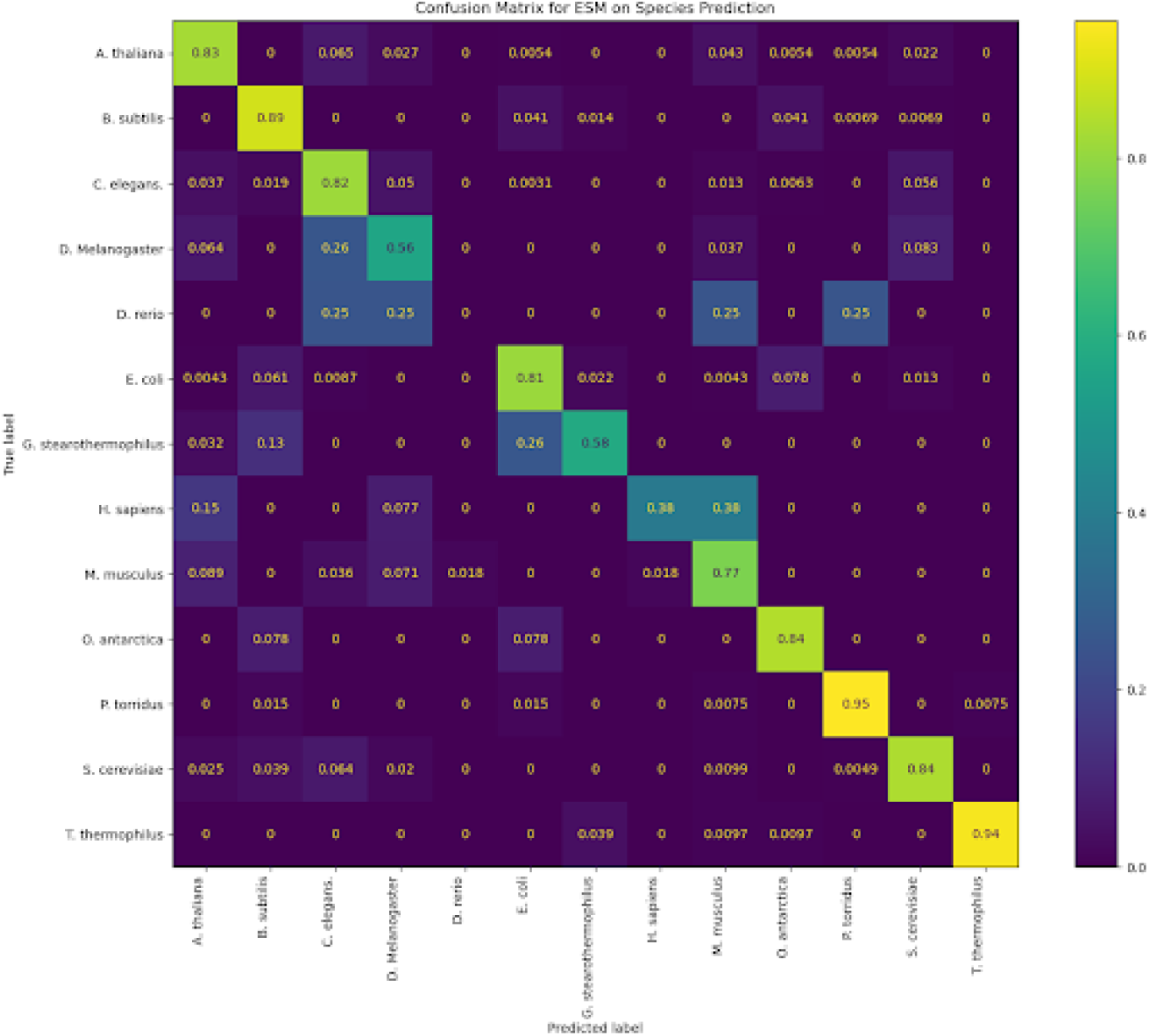
Confusion matrix for ESM model on species prediction.

**Supplementary Figure 7.**
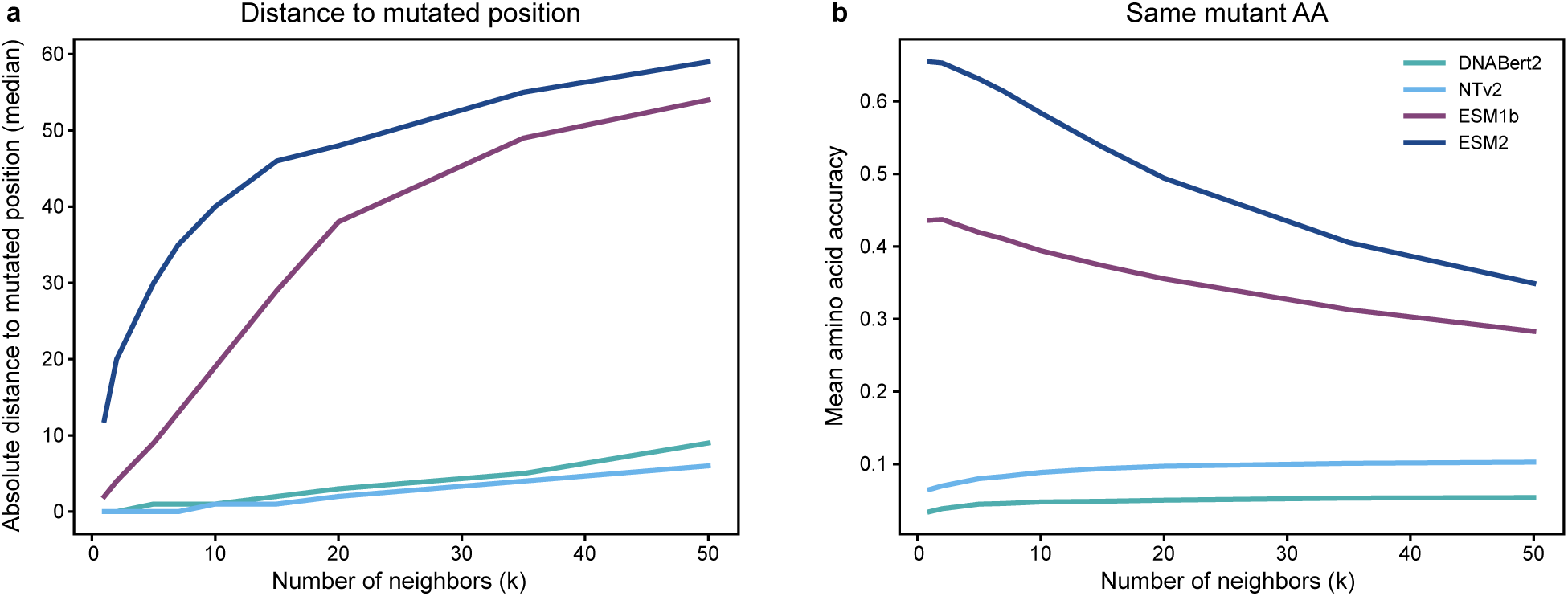
gLM embeddings capture the mutated position while pLM embeddings capture the mutant amino acid identity. **a)** Absolute distance to median mutated position of neighbouring sequences in function of the number of neighbours of each query sequence. This relationship is shown for the embeddings of the four different models. **b)** Mean accuracy for the prediction of the mutated amino acid identity of the query sequence based on the mutated amino acid of the neighbouring sequences, in function of the number of neighbours of each query sequence. This relationship is shown for the embeddings of the four different models.

## Supplementary Tables

**Table 1.** Overview of the tasks in the curated CDS dataset. Samples in each task’s dataset contain protein sequences paired with CDS sequences. Total sampled over all 3 test sets is provided for SSP.

| Dataset | Task Type | Experiment Type | # Train Samples | # Validation Samples | # Test Samples<br>Length (bp) | Mean Sequence |
| --- | --- | --- | --- | --- | --- | --- |
| Fluorescence | Regression | Single Protein Mutagenesis | 21464 | 5366 | 27217 | 714 |
| Beta-Lactamase Activity (Unique) | Regression | Single Protein Mutagenesis | 3457 | 865 | 1080 | 858 |
| Beta-Lactamase Activity (Complete) | Regression | Single Protein Mutagenesis | 11252 | 2814 | 1080 | 858 |
| Stability | Regression | Multi-Protein | 53700 | 2512 | 12851 | 135 |
| Melting Point | Regression | Multi-Protein | 9432 | 1064 | 1648 | 1176 |
| Secondary Structure Prediction | Per AA Classification | Multi-Protein | 6224 | 1556 | 334 | 724 |

**Table 2.**
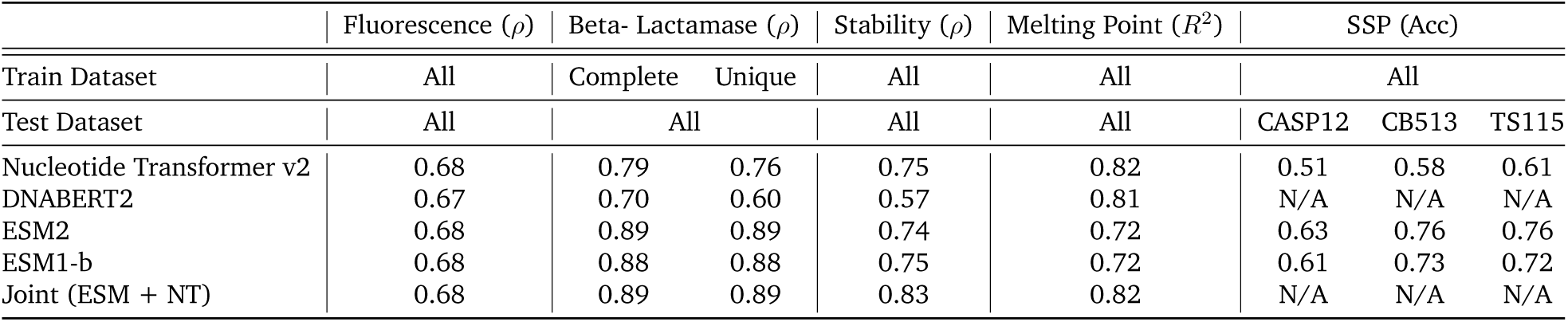
Evaluation results of Nucleotide Transformer v2 (500M), DNABERT2, ESM2 (650M), ESM1-b (650M), and joint ESM-NTv2 models on the test sets of the different tasks. The metrics used to measure performance in each task were chosen to match previous benchmarks and include Spearman correlation *ρ*, *R*^2^, and accuracy, with a higher value indicating better performance for all metrics. We note that protein models evaluated on *Complete* splits see identical proteins with different labels, but we still performed the evaluation for completeness. DNABERT2 was not evaluated on SSP since its BPE tokenization prevents residue-level predictions.

**Table 3.**
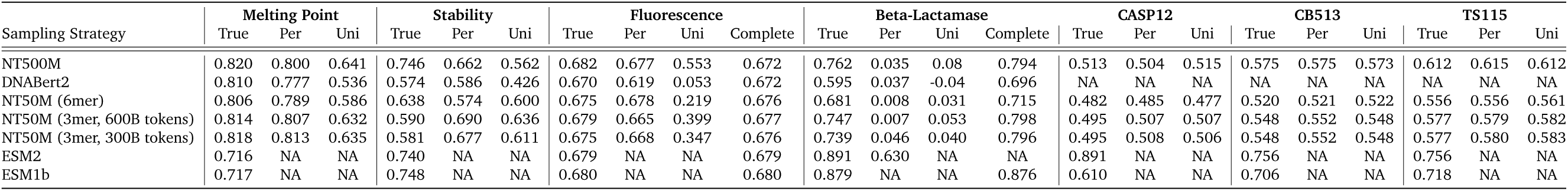
Performance of the different models across all tasks using different CDS datasets based on different codon sampling strategies (True CDS, Permutated, Unique and Complete).

**Table 4.**
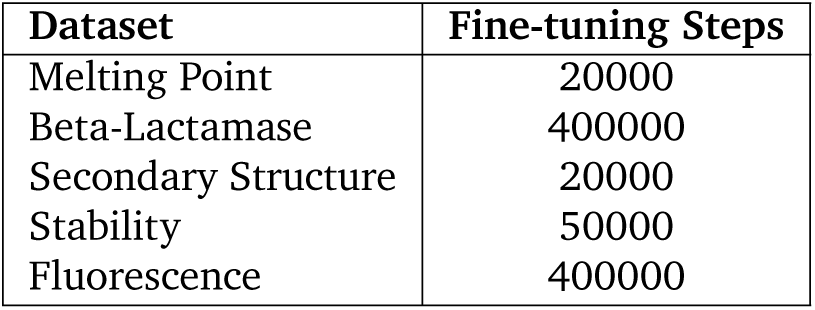
Number of Fine-tuning Steps per Dataset.

| Dataset | Fine-tuning Steps |
| --- | --- |
| Melting Point | 20000 |
| Beta-Lactamase | 400000 |
| Secondary Structure | 20000 |
| Stability | 50000 |
| Fluorescence | 400000 |

**Table 5.** Performance of NTv2 and ESM models on the melting point prediction task when trained solely on protein/CDS sequences (Nothing) or with additional information about GC-content, species of origin, or both. The metric used is the R2.

| Appended | Nothing | GC | Species | GC + Species |
| --- | --- | --- | --- | --- |
| ESM2 | 0.716 | 0.741 | 0.787 | 0.793 |
| NTv2 | 0.820 | 0.820 | 0.820 | 0.821 |

**Table 6.** Breakdown of Secondary Structure Types.

| Helices |  |  | Strands |  | Coils |  |  |
| --- | --- | --- | --- | --- | --- | --- | --- |
| helix | π-helix | α-helix | β-bridge | β-sheet | turn | bend | coil |

